# Bayesian Differential Analysis of Cell Type Proportions

**DOI:** 10.1101/2023.01.17.524410

**Authors:** Tanya T. Karagiannis, Stefano Monti, Paola Sebastiani

**Affiliations:** Institute for Clinical Research and Health Policy Studies, Tufts Medical Center, Boston, MA, USA; Division of Computational Biomedicine, Boston University School of Medicine, Boston, MA, USA; Department of Biostatistics, Boston University School of Public Health, Boston, MA, USA; Bioinformatics Program, Boston University, Boston, MA, USA

**Keywords:** Single cell transcriptomic, cell type composition, Bayesian multinomial regression

## Abstract

The analysis of cell type proportions in a biological sample should account for the compositional nature of the data but most analyses ignore this characteristic with the risk of producing misleading conclusions. The recent method scCODA appropriately incorporates these constraints by using a Bayesian Multinomial-Dirichlet model that requires a reference cell type to normalize the distribution of all cell types. However, a reference cell type that is stable across biological conditions may not always be available. Here, we present an approach that uses a Bayesian multinomial regression for the analysis of single cell distribution data without the need for a reference cell type. We show an implementation example using the rjags package within the R software.

## 1. INTRODUCTION

Single cell transcriptomics allows us to explore changes of cell type composition across conditions (Luecken and Theis 2019). Most methods analyze the changes of the proportion of each cell type across biological conditions independently of the other cell types, when in fact the cell type proportions within a sample are dependent on each other and constrained to sum to 1 (Haber *and others* 2017; Luecken and Theis 2019; Hashimoto *and others* 2019; Wilk *and others* 2020; Zhu *and others* 2020; Zheng *and others* 2020). The recent method scCODA, appropriately accounts for these constraints by using a Bayesian Multinomial-Dirichlet model (Büttner *and others* 2021). scCODA uses a multinomial distribution to describe the vector of probabilities (proportions) of all cell types in a sample, and a logit-type parameterization that relies on a reference cell type to avoid issues of convergence of the Bayesian estimation algorithm. This approach is ideal when one can identify a reference cell type whose proportion is unaffected by the condition under study and/or is stable in relative abundance across samples. However, there are situations where no such reference cell type can be determined. For example, in our ongoing study of the distribution of peripheral blood mononuclear cells (PBMCs) with age, we could not identify a cell type with stable proportion in various age groups (Karagiannis *and others* 2022). To circumvent this problem, we present an alternative Bayesian multinomial regression analysis of cell type composition. The approach estimates the cell type proportions without the need to provide a reference cell type. We describe the approach and provide an example analysis script in the R software.

## 2. METHODS

### 2.1 Modeling approach

We have configured the Bayesian multinomial regression using the R package *rjags* (Plummer 2008) to model the cell type abundance distribution as a function of covariates of interest. We model the vector of cell type counts in a sample using the following parameterization:

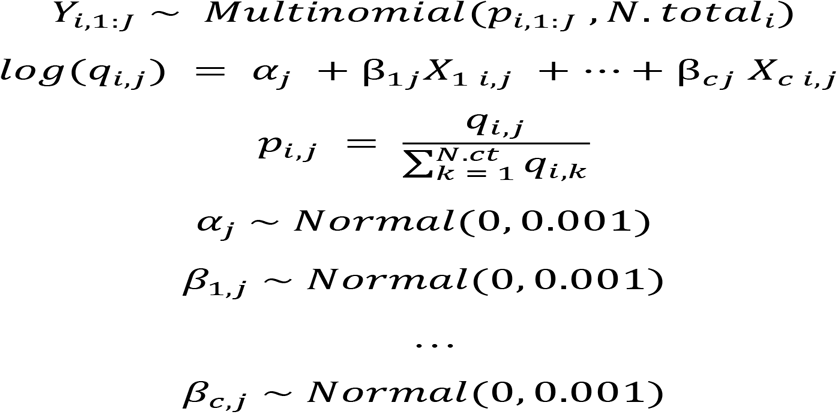

where *Y*_*i*,1:*J*_ represents the vector of numbers of cell types 1: *J* in sample *i*, and is modeled using a multinomial distribution with probabilities *p*_*i*,1:*J*_ such that 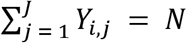. *total*_*i*_ and 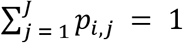, for all sample *i*. The probabilities *p*_*i*,1:*J*_ can depend on covariates *X*_1_… *X*_*c*_ through the function *log*(*q*_*i*,*j*_). The regression parameters *α*_*j*_, β_1*j*_,…, β_*cj*_, *j* = 1: *J* can be estimated using Markov Chain Monte Carlo (MCMC) sampling as implemented in rjags (Plummer 2008), and used to estimate the probabilities of cell types.

The advantage of this Bayesian and unconditional approach is that one can use many tools to monitor the goodness of fit of the model and the convergence of the parameter estimates. In addition, one can estimate the absolute proportion of each cell type and provide measures of the uncertainty of the estimates. To assess the effect of covariates in each cell type, we can use the MCMC estimates of the regression coefficients and their standard errors to calculate approximate two-sided p-values and use Benjamin-Hochberg correction for multiple testing. In addition, the analysis produces estimates of the absolute proportions of cell type per sample that are easier to interpret compared to odds ratios.

### 2.2 Analysis script

We developed an example analysis script that uses this approach in the R packages rjags (Plummer 2008) and coda (Plummer *and others* 2006). The script can be easily adjusted based on the study design and covariates of interest (Figure S1). To run the analysis scripts, the program JAGS (https://mcmc-jags.sourceforge.io/) is required for download and installation. JAGS is a program for statistical analysis in the Bayesian framework using MCMC simulations. To run the analysis scripts for model configuration, initialization and parameter inference, the R packages rjags (Plummer 2008) and coda (Plummer *and others* 2006) are required for installation. Additional R packages required for data initialization, manipulation, and visualization include packages in tidyverse, and the hablar and patchwork packages.

### 2.3 Data

We demonstrate this model and approach using cell type distribution data of 66 subjects from single cell transcriptomics datasets of aging and longevity. The data is described in Karagiannis et al (Karagiannis *and others* 2022).

## 3. APPLICATION

As an example, we used this approach to characterize the distribution of PBMCs at different ages. We used single cell transcriptomics data of PBMCs from 66 male and female subjects across four age groups with ages 20-119 years to identify 13 immune cell types based on specific gene signatures (Karagiannis *and others* 2022). We applied the proposed Bayesian multinomial regression model to the distributions of the 13 immune cell types and used 1,000 MCMC iterations with 500 iterations for burn-in to estimate cell type proportions and 95 percent credible intervals for males and female subjects for each age group across all immune cell types.

Figure 1 displays the estimates of the 13 cell types using this approach and the observed cell type proportions calculated in the 66 subjects grouped by age and sex. The plots show a very good agreement between observed and estimated proportions, particularly for not small probability values. This analysis identified significant age-related changes of cell type composition in EL including a significant reduction of lymphocyte subtypes nCD4TC and mCD4TC (females: 6.00-9.34%; males: 5.38-7.78%) compared to younger age (females: 21.62-32.10%; males: 18.18-29.08%) and a significant decrease of mDC and pDC in EL (females: 0.31-0.70%; males: 0.32-0.88%) compared to younger age (females: 0.80-1.05%; males: 0.82-1.33%). Comparing the estimated cell type proportions to the relative proportions across subjects, we found similar results across cell types. Full results are described in Karagiannis et al (Karagiannis *and others* 2022).

**Figure 1.**
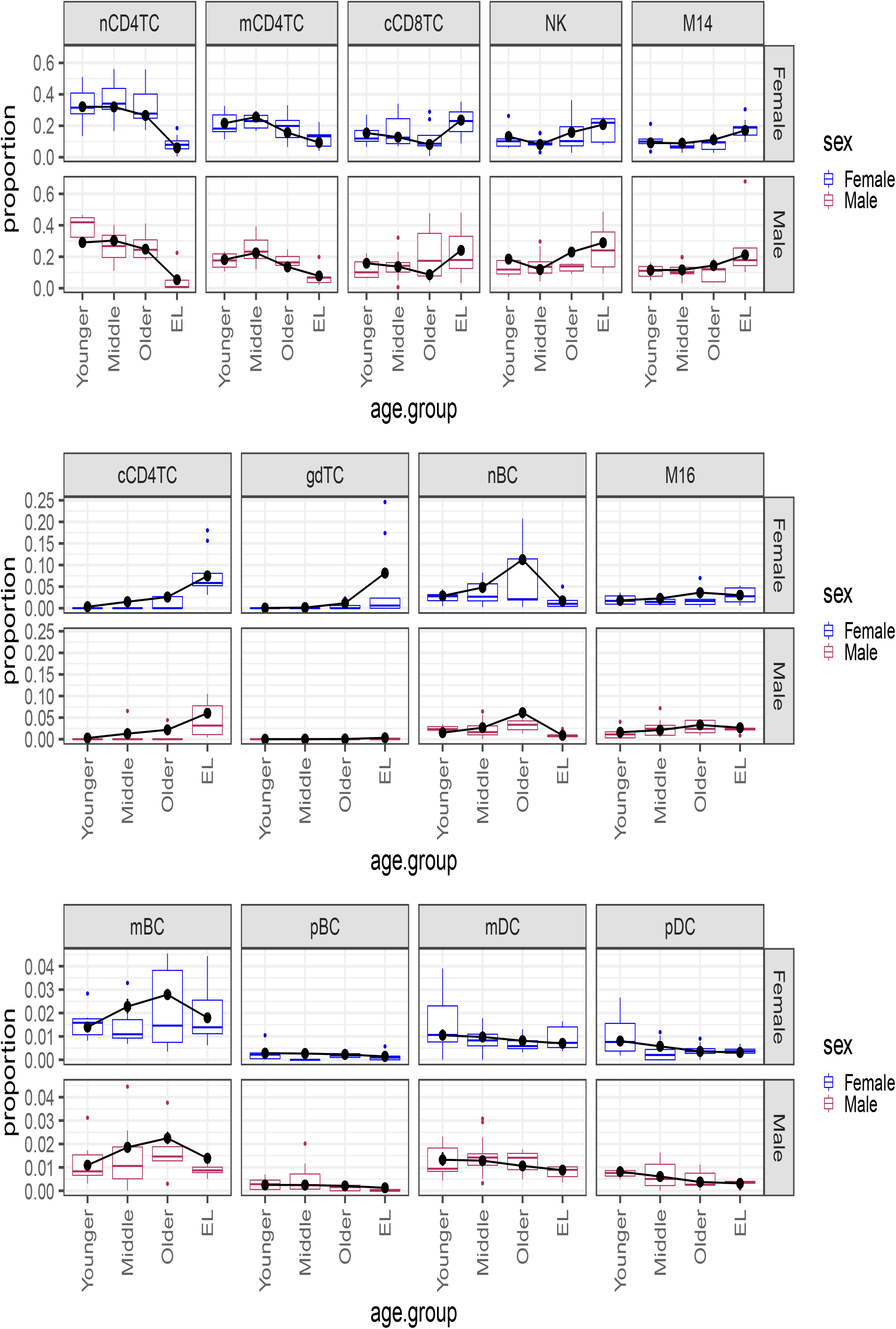
Multinomial regression cell type composition estimates across age and sex in PBMCs. Plot of the Bayesian estimates and observed relative proportions of the 13 immune cell types in PBMCs in each age group (Younger, Middle, Older, EL), for males and females. We applied the Bayesian multinomial regression to a matrix of the13 cell counts across the 66 subjects. The 13 cell types are: noncytotoxic naive and memory CD4^+^ T cells (nCD4TC, mCD4TC), cytotoxic CD4^+^ T cells (cCD4TC), cytotoxic CD8^+^ T cells (cCD8TC), gamma-delta T cells (gdTC), naive, memory and plasma B cells (nBC, mBC, and pBC), Natural Killer cells (NK), CD14^+^ and CD16^+^ monocytes (M14 and M16), and myeloid and plasmacytoid dendritic cells (mDC and pDC). The relative proportions per subject are represented as boxplots for Females (blue) and Males (maroon). For each cell type, the estimates are overlayed with points (black) and connected by a line (black) to highlight trends across age groups.

For comparison, we applied scCODA to the distribution of the 13 immune cell types. When no obvious reference cell type is available, the recommended use of scCODA is to run the analysis using each cell type as a reference, for a total of 13 tests for comparison in our case, and to then call as significant those changes observed with a credible effect in more than 50% of the runs. Table 1 displays the credible compositional changes between EL and younger age for each cell type identified by the Bayesian multinomial regression and by scCODA. We found that scCODA identified compositional changes in 4 of the 9 cell types identified as significantly changed by the multinomial regression model. Of note, although scCODA found nBC to have a credible change in EL compared to younger age, it only identified this change in 10 out of the 13 tests run. The decrease in composition of nBC in centenarians has been previously reported (Hashimoto *and others* 2019) and we were able to confirm this credible decrease using the multinomial regression model. We also identified a significant increase in M14 composition using the multinomial regression that supports previous reports of increased composition of M14 with age (Zheng *and others* 2020). However, scCODA only identified the credible change in M14 in 2 out of the 13 tests run.

**Table 1.** Table of significant cell type specific credible changes in composition between EL and younger age based on the Bayesian multinomial regression and scCODA.

| cell types | scCODA: number of credible changes | scCODA: percent of credible changes | scCODA: credible effect | multinomial: credible effect |
| --- | --- | --- | --- | --- |
| nCD4TC | 12 | 0.923077 | TRUE | TRUE |
| mCD4TC | 11 | 0.846154 | TRUE | TRUE |
| cCD4TC | 12 | 0.923077 | TRUE | TRUE |
| cCD8TC | 2 | 0.153846 | FALSE | FALSE |
| gdTC | 0 | 0 | FALSE | TRUE |
| nBC | 10 | 0.769231 | TRUE | TRUE |
| mBC | 0 | 0 | FALSE | FALSE |
| pBC | 0 | 0 | FALSE | TRUE |
| NK | 2 | 0.153846 | FALSE | FALSE |
| M14 | 2 | 0.153846 | FALSE | TRUE |
| M16 | 0 | 0 | FALSE | FALSE |
| mDC | 0 | 0 | FALSE | TRUE |
| pDC | NaN | NaN | FALSE | TRUE |

In summary, application of our Bayesian multinomial regression model identified multiple age-related changes including those previously reported and showed that this method can be used to identify and provide simple interpretations of changes in the distribution of cell types without a reference cell type.

## DISCUSSION

We have presented an implementation of Bayesian multinomial regression to analyze single cell distribution data that accounts for cell type proportion compositional constraints within each sample and does not require the choice of a reference cell type. The analysis script we developed uses the rjags package in the R software and can be easily generalized to different sets of covariates and study design. We provide detailed documentation for model and parameter configuration and initialization as well as for application to single cell distribution data to obtain posterior distributions of sample proportions across conditions. As shown in the application to distributions of PBMCs with age, the Bayesian multinomial regression allows for the investigation of cell type specific compositional changes applicable to studying disease and other conditions. An important feature of our unconditional approach is to estimate the absolute proportions of cell type per sample that are easier to interpret compared to odds ratios or other relative metrics.

## Software

The model and application scripts in the R software are available on github (https://github.com/Integrative-Longevity-Omics/Bayesian-Multinomial-Regression).

## Supplementary Material

Supplementary material is available online at http://biostatistics.oxfordjournals.org.

## Acknowledgements

TK, SM, PS are supported by NIH-NIA UH2AG064704.

