## Supplementary material for "Bayesian Differential Analysis of Cell Type Proportions": Figures S1

**Supplementary Materials**

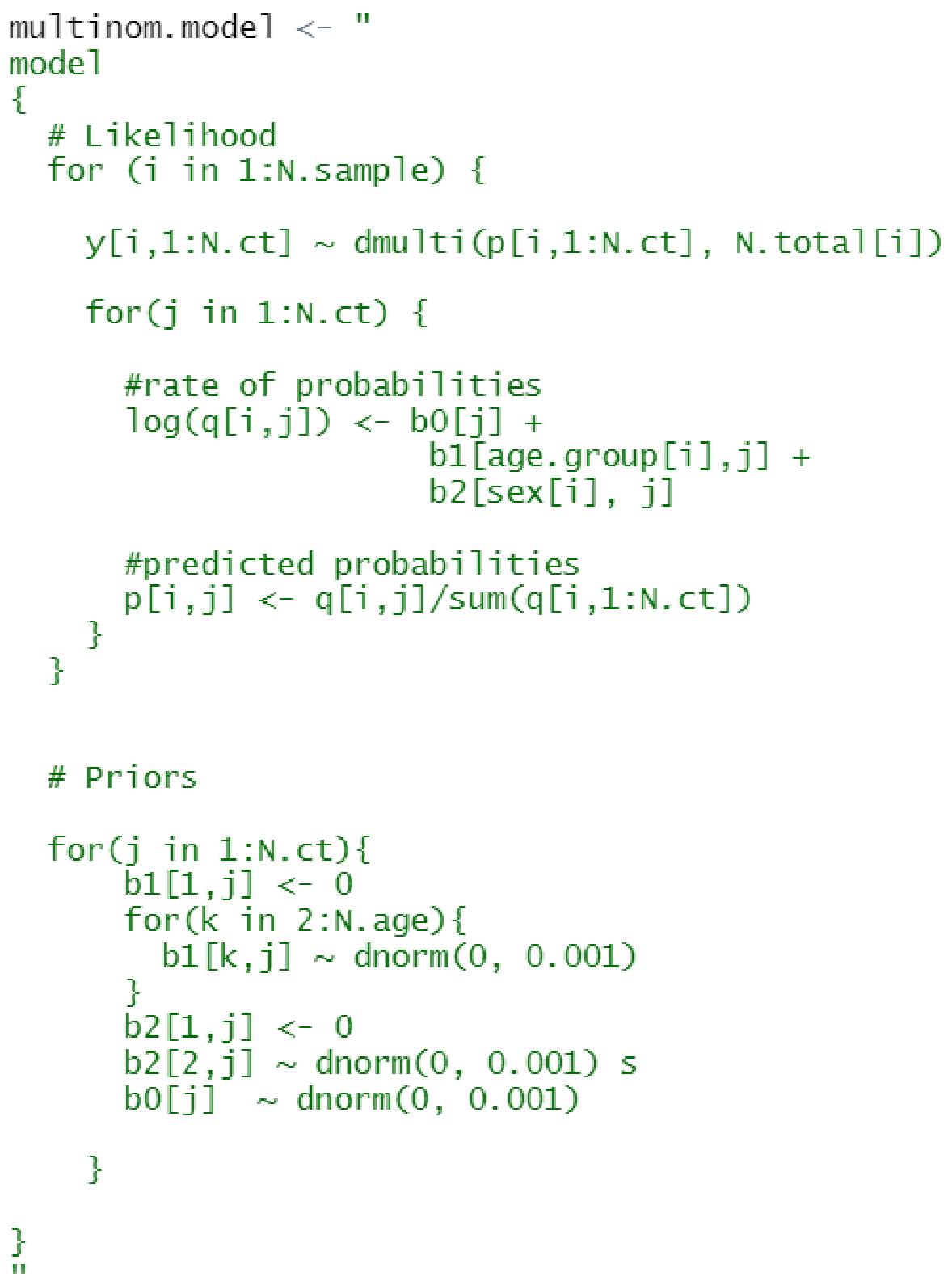

**Figure S1. Bayesian multinomial regression model configuration using rjags package in the R software.**
